# High-resolution *de novo* structure prediction from primary sequence

**DOI:** 10.1101/2022.07.21.500999

**Authors:** Ruidong Wu, Fan Ding, Rui Wang, Rui Shen, Xiwen Zhang, Shitong Luo, Chenpeng Su, Zuofan Wu, Qi Xie, Bonnie Berger, Jianzhu Ma, Jian Peng

## Abstract

Recent breakthroughs have used deep learning to exploit evolutionary information in multiple sequence alignments (MSAs) to accurately predict protein structures. However, MSAs of homologous proteins are not always available, such as with orphan proteins or fast-evolving proteins like antibodies, and a protein typically folds in a natural setting from its primary amino acid sequence into its three-dimensional structure, suggesting that evolutionary information and MSAs should not be necessary to predict a protein’s folded form. Here, we introduce OmegaFold, the first computational method to successfully predict high-resolution protein structure from a single primary sequence alone. Using a new combination of a protein language model that allows us to make predictions from single sequences and a geometry-inspired transformer model trained on protein structures, OmegaFold outperforms RoseTTAFold and achieves similar prediction accuracy to AlphaFold2 on recently released structures. OmegaFold enables accurate predictions on orphan proteins that do not belong to any functionally characterized protein family and antibodies that tend to have noisy MSAs due to fast evolution. Our study fills a much-encountered gap in structure prediction and brings us a step closer to understanding protein folding in nature.

## Main Text

Half a century after Anfinsen demonstrated a connection between a protein’s amino acid sequence and folded three-dimensional conformation, scientists have built deep learning models that finally can predict high-resolution protein structures. The recent success of DeepMind’s AlphaFold2 (*1*) and RoseTTAFold (*2*) in protein structure prediction has mainly been based on advances in deep learning, especially transformer-based models (*3*), and the accumulation of large databases of protein sequences and structures that enable effective training of large models. Both these methods need as input evolutionary data in the form of multiple sequence alignments (MSAs) of homologous sequences *aligned* to the primary one, a technique which has been a staple of structure prediction methods (*1, 2, 4–15*). By extracting residue-residue covariances from these MSAs, these algorithms have been shown to greatly outperform previous approaches, including physics-based models, homology-based methods and convolutional neural networks (*16*) and predict structures with atomic-level accuracy for the first time in history. Furthermore, many methods have since built upon AlphaFold2 for other prediction tasks in structural biology, including protein-protein interactions, disordered regions, and binding sites (*17–20*). However, prediction accuracies for all these advanced methods drop sharply in the absence of a multitude of sequence homologs from which to construct MSAs.

Instead, we sought to train an algorithm that learns to model protein three-dimensional structure without relying on MSA preprocessing (i.e., it is alignment-free). We hypothesize that akin to extracting grammatical structure from large collections of natural language corpuses, predicting protein structure from protein sequence databases should be possible without having to rely on aligned MSAs. The use of deep language models has recently led to major progress in language modeling (*21*) as well as elucidating protein properties (*22–25*), in protein-protein interaction prediction (*26, 27*), and in viral escape prediction (*28*) from sequence data. We reasoned that the transformer’s attention mechanism used to model long-range relationships in natural language sequences should also be applicable to extracting correlations from evolutionary relationships present in protein sequences (**Fig 1A**). In contrast to natural languages, proteins are more than merely strings; they are physical chains of amino acids that fold into three-dimensional structures. Thus, to model 3D protein structures, we were motivated to incorporate geometric intuition into the transformer architecture design.

**Fig. 1.**
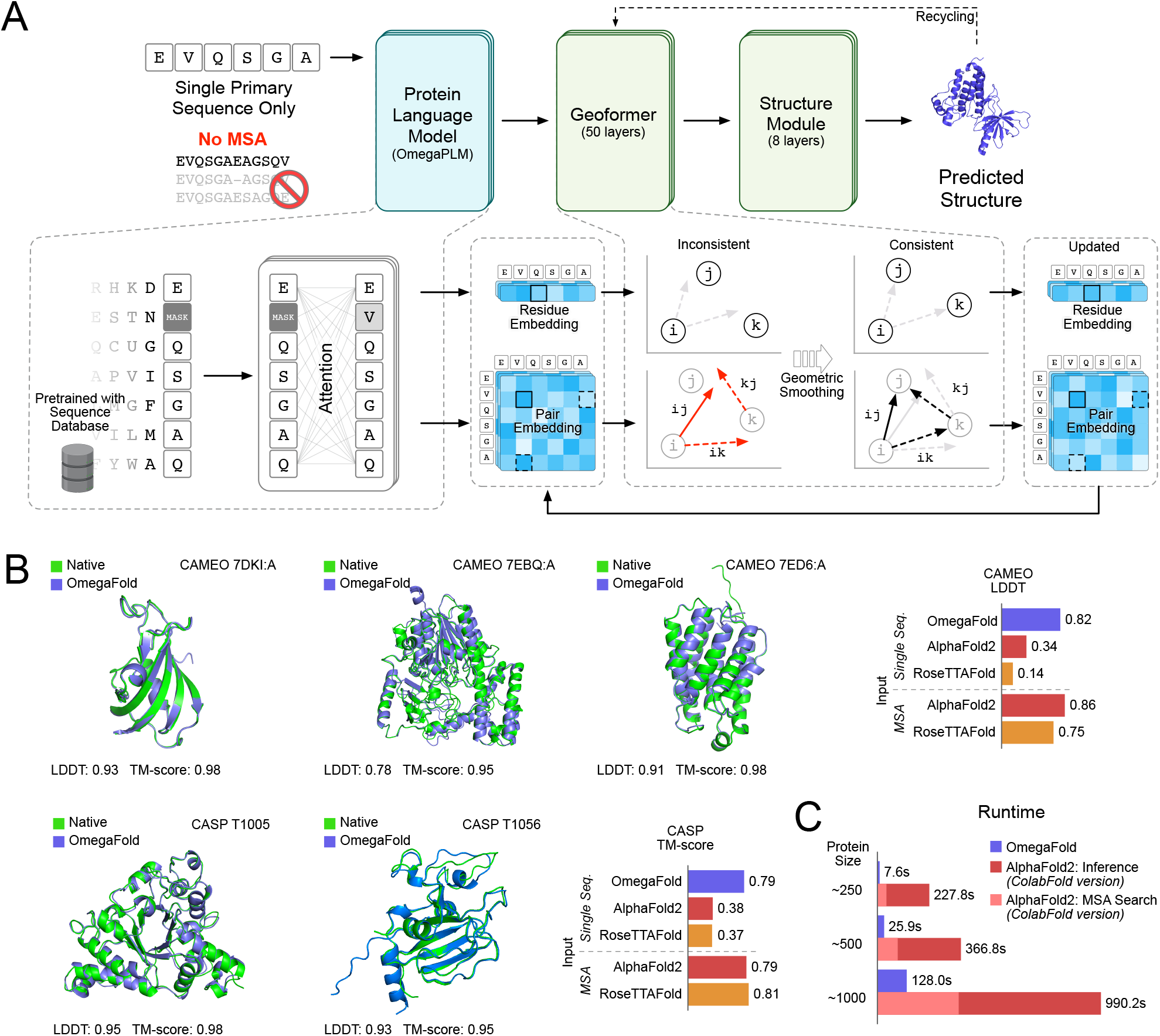
Overview of OmegaFold and Results. **(A)** Model architecture of OmegaFold. The primary protein sequence is first fed into a pretrained protein language model (OmegaPLM) to obtain residue-level node embeddings and residue-residue pairwise embeddings. A stack of Geoformer layers then iteratively updates these embeddings to improve their geometric consistency. Lastly, a structure module predicts the 3D protein structures from the final embeddings. The predicted structure and the embeddings could be fed again as input into another cycle through a recycling procedure to predict a more refined structure. **(B)** Evaluations on recent CAMEO an CASP targets. Our predictions (blue) for 7DKI:A, 7EBQ:A, 7ED6:A from CAMEO and T1005, T1056 from CASP are highly accurate according to the experimental structures (green). Figures on the right show held-out test results on 146 CAMEO targets and 29 challenging CASP targets. OmegaFold significantly outperforms AlphaFold2 and RoseTTAFold when only single sequences are provided as input on both standard CAMEO Local Distance Difference Tests (LDDTs) and CASP TM-scores; OmegaFold performs comparably to AlphaFold2 and RoseTTAFold on the CASP and CAMEO test cases when the standard MSAs are used as input. **(C)** Runtime analysis. OmegaFold is significantly faster than AlphaFold2 (ColabFold version) on single-chain proteins with typical lengths of around 250, 500 and 1000 residues. ColabFold was used to further decrease the runtimes of the MSA search time (pink) and model inference time (red).

We propose OmegaFold, to our knowledge the first computational method to predict the structure of a protein from its primary sequence alone with high accuracy, using a new combination of a large pretrained language model for sequence modeling and a geometry-inspired transformer model for structure prediction (**Fig. 1A**). Notably, OmegaFold requires only a single amino-acid sequence for protein structure prediction, does not rely on MSAs nor known structures as templates, and scales roughly ten-times faster with comparable or better accuracy to MSA-based methods such as AlphaFold2 and RoseTTAFold. We demonstrate OmegaFold’s ability to more accurately predict the structures of orphan proteins and antibodies, for which evolutionary information is scarce or noisy, respectively (**Fig. 2**).

**Fig. 2.**
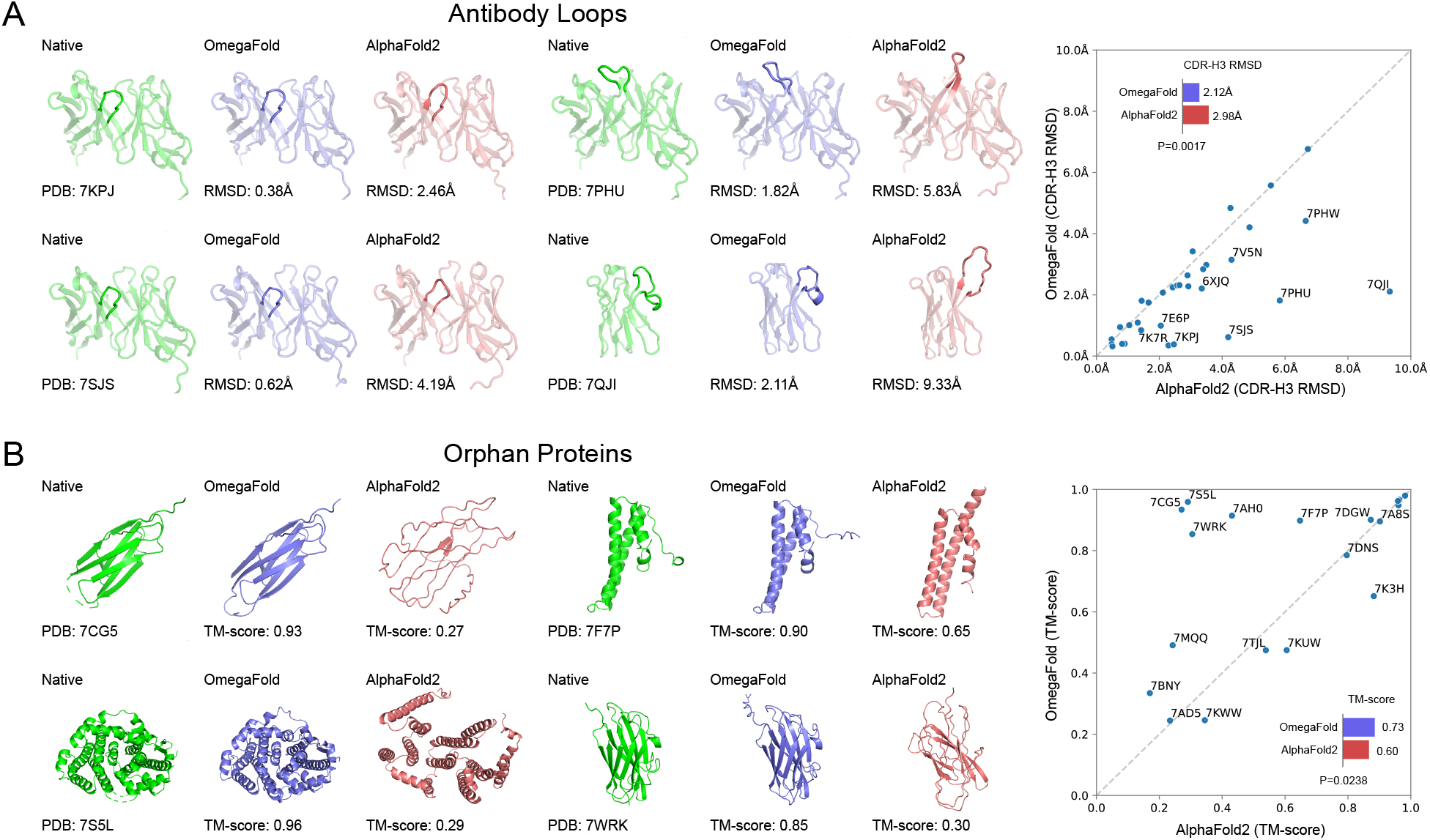
OmegaFold performs significantly better than AlphaFold2 in modeling antibody CDR H3 regions and predicting structures of orphan proteins. **(A)** Antibody CDRH3 regions. OmegaFold predictions are colored blue; AlphaFold2 predictions,red; and native experimental structures, green. CDR H3 regions are highlighted in plots. On CDR3 of nanobody 7QJI and CDRH3 loops of antibodies 7KPJ, 7PHU, 7SJS, root mean square distances (RMSDs) of OmegaFold loop predictions are significantly lower than AlphaFold2 predictions. The scatter plot depicts the comparison on 33 recently released nanobody and antibody proteins with highresolution experimental structures. Overall, OmegaFold predictions (RMSD=2.12Å) are significantly better than AlphaFold2 predictions (RMSD=2.98 Å), with a P-value of 0.0017. **(B)** Orphan proteins. On orphan proteins 7CG5 (sensor domain of RsgI4), 7F7P (anti-CRISPR, AcrIIC4), 7S5L (Cembrene A synthase) and 7WRK (hypothetical protein TTHA1873), TM-scores—a common metric for assessing the topological similarity of protein structures—of OmegaFold predictions are significantly higher than AlphaFold2 predictions. The scatter plot shows comparisons on 19 recently released orphan proteins with no homologous sequences identified. Overall, OmegaFold predictions (TM-score=0.73) are better than AlphaFold2 predictions (TM-score=0.60), with a P-value of 0.0238.

### The OmegaFold model

The overall model of OmegaFold is conceptually inspired by recent advances in language models for natural language processing coupled with deep neural networks used in AlphaFold2. We introduce OmegaPLM, a deep transformer-based protein language model (PLM), trained on a large collection of unaligned and unlabeled protein sequences, to learn single- and pairwise-residue embeddings (or representations) as powerful features that model the distribution of sequences; OmegaPLM is able to capture structural and functional information encoded in the amino-acid sequences through the embeddings. These are then fed into Geoformer, a new geometry-inspired transformer neural network, to further distill the structural and physical pairwise relationships between amino acids. Lastly, a structural module predicts the 3D coordinates of all heavy atoms (**Fig. 1A**; **Materials and Methods**).

The key idea behind Geoformer is to make the embeddings from our language model more geometrically consistent—amino acid node and pairwise embeddings generate consistent coordinates and distance predictions when projected to 3D. While similar in principle to the Evoformer module in AlphaFold2 which applied attention mechanisms for information integration, ours mainly focuses on vector geometry as opposed to evolutionary variation. It consists of a deep stack of 50 Geoformer layers, which is inspired by the fundamental theorem of vector calculus in geometry (**Fig. 3B**). Each Geoformer layer encodes information in node representations (*s*_*i*_) for residue *i* and pairwise representations (*p*_*ij*_) between residues *i* and *j* by enforcing their geometric consistency. Intuitively, we can view the representation of residue *i* in a high-dimensional vector space, and each pairwise representation *p*_*ij*_ is a vector pointing from residue *i* to residue *j*, which will be used for predicting three-dimensional coordinates and distances. Ideally, coordinates and pairwise distances predicted from *s*_*i*_ and *p*_*ij*_ should satisfy properties of Euclidean geometry, such as triangular inequality. However, these properties may not always hold if the representation vectors and predictions are output from neural networks. Based on these geometric insights, we designed a geometry-inspired nonlinear attention module to sequentially update *s*_*i*_ and *p*_*ij*_, attempting to make them consistent. By stacking Geoformer layers upon the PLM, OmegaFold captures the geometry of a protein structure with representations which are then projected onto 3D space with an 8-layer structure module (**Fig. 1A**).

**Fig. 3.**
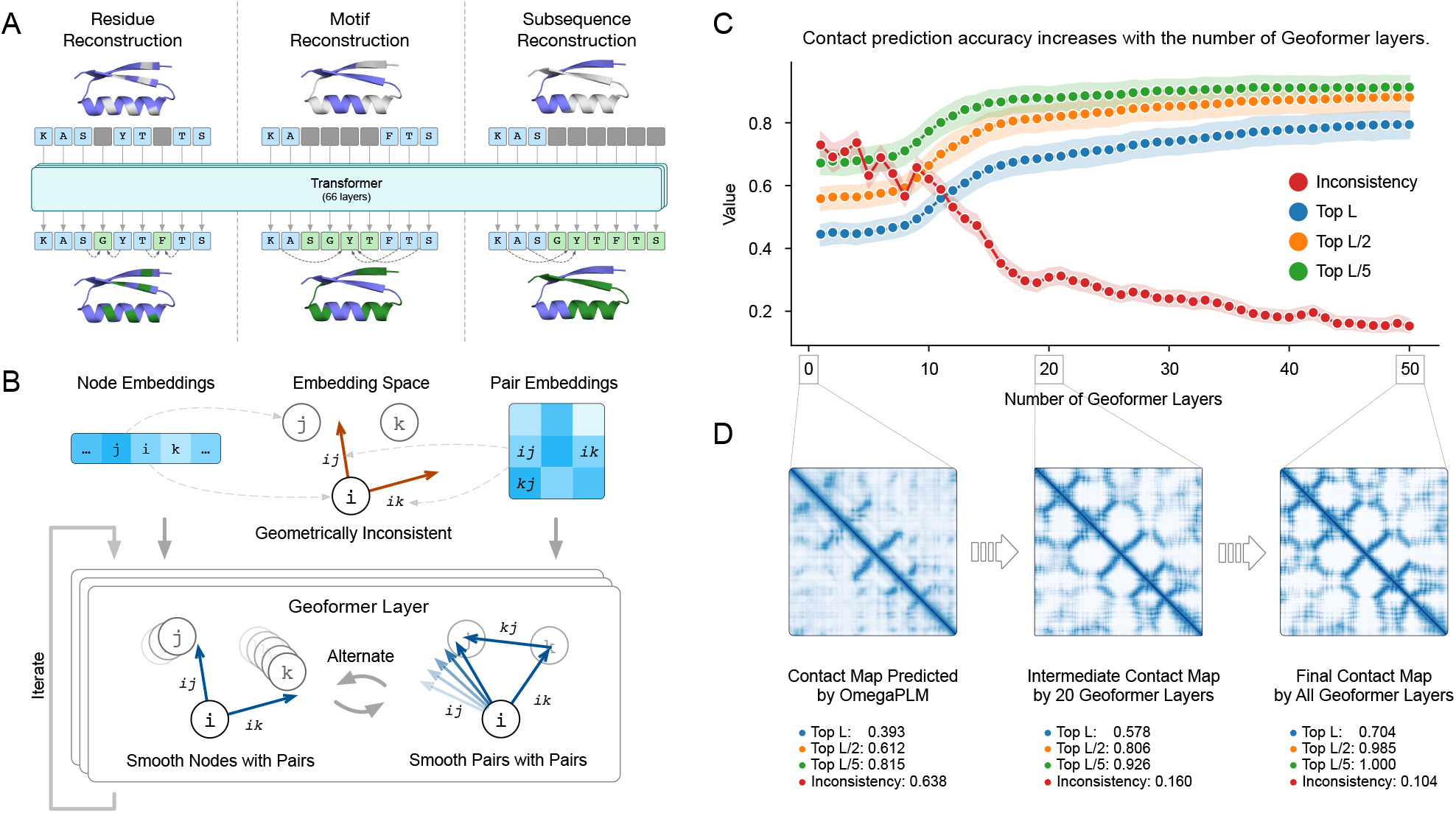
Geoformer refines the contact map predicted by protein language model (OmegaPLM) through geometric smoothing. **(A)** OmegaPLM model is pretrained by per-residue mask loss, per-motif mask loss and subsequence mask loss on unaligned protein sequences. **(B)** Geoformer layers iteratively smooth node and pairwise embeddings and reduce geometric inconsistency among them. Initially, node and pairwise embeddings generated by OmegaPLM reside in a latent space with geometric inconsistency (red). In each Geoformer layer, these embeddings are updated iteratively to refine the geometric inconsistency: each node embedding was updated with related pairwise embeddings, and each pairwise embedding was updated by triangular consistency of pairwise embeddings. **(C)** Geoformer layers improve geometry of contact predictions. Top L, Top L/2, Top L/5 accuracies denote contact prediction precision among top predicted contact pairs. Inconsistency is defined as the percentage of predicted distance triples {ij, jk, ik} that violate the triangular inequality. It is clear to see that the prediction accuracy improves while inconsistency drops with stacked Geoformer layers. **(D)** Visualization of contact maps. The contact maps predicted in the initial, 20^th^ and last Geoformer layer for protein 7EAD_A from the CAMEO dataset. In the first 20 layers, Geoformer mainly focuses on resolving the triangular inconsistency, which decreases from 0.638 to 0.160. In the last 30 layers, Geoformer mainly refines the details of contact prediction, with TopL accuracy increasing from 0.578 to 0.704.

Intrigued by AlphaFold2’s training, we trained the entire OmegaFold model, with several structural objectives, including contact prediction, Frame Aligned Point Error (FAPE) loss (*1*)—a geometrically restricted version of root-mean-squared deviation (RMSD) of relative atomic positions—and torsion angle prediction. The full model was jointly trained on ∼110,000 single-chain structures from the Protein Databank PDB (*29–31*) deposited before 2021 and all single domains from the SCOP v1.75 database with at most 40% sequence identity cutoff (*32–34*). We used later-released structures for validation and hyperparameter selection. We also excluded protein structures that appeared in our test sets as well as their homologs up to 40% sequence identity during training.

### OmegaPLM outperforms vanilla language models

As a precursory assessment of our language module, we applied three strategies for masking out residues and trained to optimize the standard Bidirectional Encoder Representations from Transformers (BERT)-style objective (*21*) to predict masked residues from the rest of the sequence using a cross-entropy loss function (**Fig. 3A**) mask out a random 15% of residues in protein sequences, 2) mask out random consecutive subsequences of 5-8 residues, and 3) mask out half of the entire protein sequence. For convenience, we chose the same hyperparameters as those used in the popular transformer network ESM-1b (*23*), including the number of layers and the number of heads but with a more lightweight implementation of the attention module (*35*). After pretraining on sequences in UniRef50 (dated at 2021/04) (*36*), our PLM demonstrates improved language modeling performance over vanilla transformer-based networks as well as contact prediction, with much smaller memory requirements (**Supplementary Figures S1 & Table S1**).

### OmegaFold performs well on CASP and CAMEO benchmark datasets

We assessed OmegaFold’s ability to predict protein structures by testing on two benchmarks: a CASP set with 29 of the most challenging proteins from the free-modeling category in two recent CASP experiments (37), and a CAMEO dataset with 146 of the most recent single-chain proteins (appearing in the first six months of the 2022 CAMEO evaluation), spanning a wide range of prediction difficulty levels. For comparison, we computed predictions as compared to AlphaFold2 and RoseTTAFold run in their default mode with MSAs as input. Remarkably, the structures predicted by OmegaFold, with a single sequence as input, were as accurate as the advanced MSA-based methods (Fig. 1B). On the CAMEO dataset, OmegaFold structures had a mean local-distance difference test (LDDT) score of 0.82, with comparable accuracy to RoseTTAFold structures (0.75 mean LDDT score) and AlphaFold2 structures (0.86 mean LDDT score) predicted from MSAs. (LDDTs are a commonly used metric for structure evaluation.) On the more challenging CASP dataset, OmegaFold structures were also quite accurate with an average TM-score—a common metric for assessing the topological similarity of protein structures—of 0.79, slightly lower than that of RoseTTAFold structures (0.81 mean TM-score) and equivalent to AlphaFold2 structures (0.79 mean TM-score).

### OmegaFold outperforms RoseTTAFold and AlphaFold2 on single-sequence inputs

We also tested OmegaFold’s relative performance using the single-sequence versions of AlphaFold2 and RoseTTAFold on these two datasets. When only a single sequence was given as input, their predicted structures were statistically highly inferior to those of OmegaFold (**Fig. 1B**), indicating that the performance of the MSA-based methods drops when evolutionary information is not given.

### OmegaFold enables accurate predictions on orphan proteins and antibodies

We further sought to assess OmegaFold’s performance in predicting challenging structures of recently released antibody and orphan proteins from the PDB, for which other methods perform poorly. Antibody complementarity-determining regions (CDRs) are the most diverse and variable parts of the molecules (*38, 39*). Because of antibodies’ fast-evolving nature, MSAs on CDRs, especially in the CDR3 loops on the heavy chain of the antibodies which—despite being highly enriched in amino acid composition— are extremely noisy. As a result, methods like AlphaFold2 are unreliable and have very low predicted LDDT (pLDDT) scores (**Fig. 2A**). Unlike antibodies, orphan proteins, by definition, lack sequence and structure homology information, and thus are also difficult to predict by MSA-based methods (**Fig. 2B, Materials and Methods**). On both antibody loops and orphan proteins, OmegaFold achieves much higher statistical prediction accuracy, in contrast to AlphaFold2, likely due to the advantages of its single sequence-based prediction method.

### Towards interpretability of OmegaFold

We sought to understand how the various components of OmegaFold contributed to prediction. To test whether Geoformer was boosting the PLM’s performance, we calculated the distance (or contact) maps using embeddings from OmegaPLM and each Geoformer layer; and computed the evolution of contact accuracy and geometric inconsistency as the number of Geoformer layers increased (**Fig. 3C & D**). We found that the contacts and distances predicted by OmegaPLM alone are reasonably accurate. Within the first 20 Geoformer layers we found that the geometric inconsistency, measured by violations of the distance triangle inequality, is greatly reduced, and the prediction accuracy of contact pairs is much improved. The remaining Geoformer layers appear to focus on reducing the geometric inconsistency until the prediction accuracy plateaus. We also demonstrated through ablation studies on component parts of OmegaFold, that the components contribute to the performance of the overall model (**Table S3**). Furthermore, when we retrained an AlphaFold2 model with only single sequences as input, we found that it was not able to match the prediction quality of OmegaFold.

### OmegaFold improves runtimes by an order of magnitude

Since MSAs are no longer required for OmegaFold to achieve high-resolution performance, its overall runtime is much faster than AlphaFold2, as well as the latest highly-optimized ColabFold-AF2 (*40*) (**Fig1. C**).

In summary, our study leverages a protein language model trained on unaligned sequences to predict protein structures from single amino acid sequences alone. We give further evidence that such evolutionary information may well be encoded in primary sequences, which can then be used as features for structure prediction. As more sequencing data accumulates to feed MSA-based methods, OmegaFold fills the interim gap, as well as importantly predicts from sequences for which MSAs are difficult to construct.

We expect that our conceptual advance and further algorithmic development along these lines will continue to enable a wide spectrum of protein science applications, such as multi-state conformational sampling, where AlphaFold2 has had some success to regions for which predictions are challenging (*41*), variant effect prediction, protein-protein interactions and protein docking.

## References and Notes

### Author contributions

J.P., J.M., B.B., F.D., R.Wu developed the conceptual ideas. R.Wu, F.D., R.Wang and R.S. implemented the main algorithms and finished model training. X.Z., R.Wu and R.Wang did data preprocessing. C.S., Z.W. and R.Wu conducted baseline experiments. B.B., J.M., J.P., C.S., R.Wang, R.Wu interpreted the results. B.B., J.M., J.P., F.D., R.Wang, R.Wu, R.S., X.Z., S.L., C.S., Z.W., Q.X. wrote the manuscript.

### Competing interests

J.M., J.P., F.D., R.Wang, R.Wu, R.S., X.Z., S.L., C.S., Z.W.are from HeliXon Ltd.

### Supplementary Materials and Methods

Materials and Methods

Figs. S1 to S2

Tables S1 to S4

## Supplementary Materials for

### 1 OmegaPLM: Pre-trained Protein Language Model

Unsupervised pre-trained models have achieved tremendous success in research areas such as natural language processing (*1–4*), computer vision (*5–8*) and single cell analysis (*9, 10*). The central idea is to first pre-train the models using large-scale unlabeled data, and then adopt the models to various downstream tasks with limited labeled data. One of the successful pre-trained models in biology is the pre-trained protein language models, which has been demonstrated to be able to extract biologically meaningful representations for various tasks such as protein contact prediction (*11–14*), protein function prediction (*15, 16*), zero-shot mutation effect prediction (*17*), and viral evolution and escape prediction (*18, 19*).

One of the key components of AlphaFold2 is to predict the interactions between amino acids which are far apart on the linear sequence by extracting the co-evolution information from Multiple Sequence Alignment (MSA) using the IPA module (*20*). However, the MSA information is not always available due to technical bias introduced by the homologous detection software such as Blast (*21*) and HHblits (*22*), which poses a new computational challenge that how to accurately predict the protein 3D structures solely based on the protein sequences without relying on the MSA information. However, current machine learning models could only extract weak co-evolution signals from the protein sequences, which is not sufficient to predict the protein 3D structures (*23*). Instead of constructing a general pre-trained protein language model (PLM) suitable for various protein tasks, in this work, we aim to design a PLM which could greatly benefit protein 3D structure prediction. OmegaPLM strikes a good balance between efficiency and performance, as shown in Figure S1 and Table S1, where OmegaPLM consumes less resources and performs better.

#### 1.1 OmegaPLM Architecture

An important observation is that the deeper layers of current transformer-based PLMs focus relatively more on learning protein contacts, whereas the secondary structure information is spread more evenly across both shallow and deep layers (*15*). More analysis of the ESM-1b model indicates that long-range protein contacts could be more accurately predicted by using the attention score from deeper layers compared to shallow layers. We hypothesize that the co-evolution information could be more efficiently represented by stacking more self-attention layers and encoding more training data. Therefore, in this work we focus on designing a memory-efficient self-attention architecture to allow the PLM to go deeper by improving different components of previous PLMs such as positional encoding function, non-linear transformation and normalization functions. Similar to the ESM-1b model (*12*), the overall architecture of OmegaPLM is a self-attention model (*24*) in which each token is a single amino acid. The OmegaFold model processes a protein sequence with a stack of GAU layers from (*25*) instaed of self-attention layers and multi-layer perceptrons. Our model contains 66 layers with around 670 million parameters without sharing parameters, which doubles the layer count of ESM-1b but roughly retains the parameter count. With ***n***_*i*_ ∈ ℝ^*d*^ as the d-dimensional vector representation of token at position i, we present OmegaPLM in Algorithm 1.

##### Algorithm 1

Protein language models based on the Gated Attention Module (GAU)

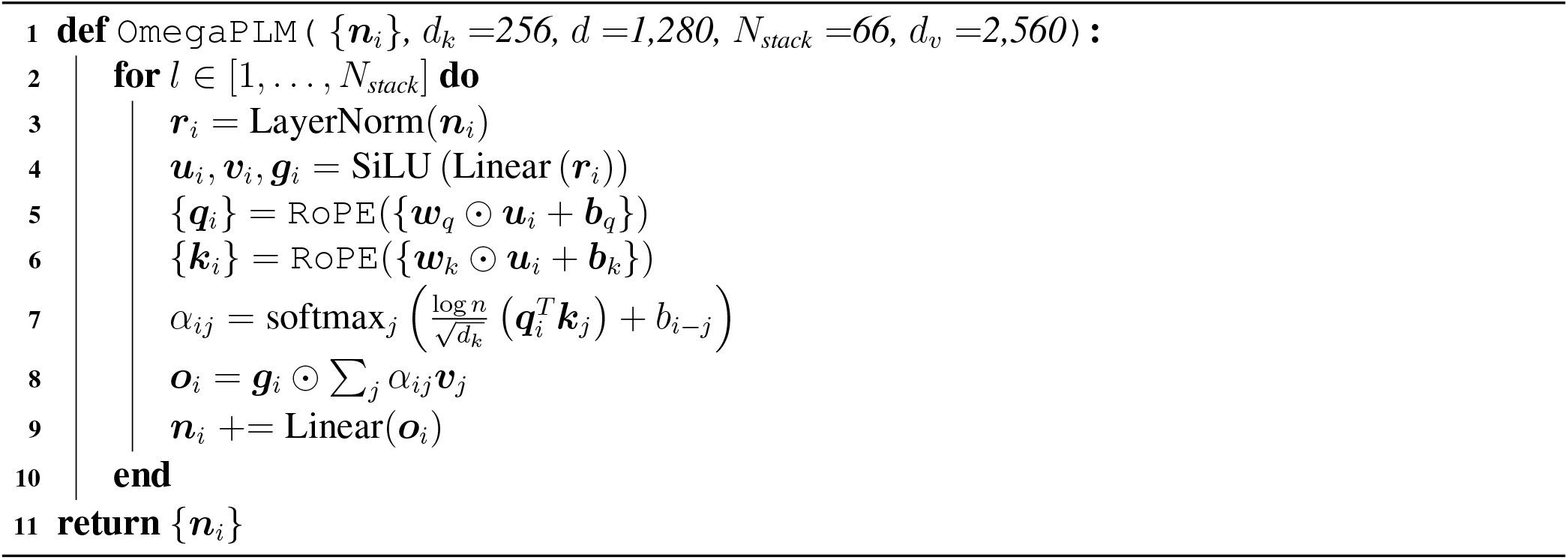

##### Pre-LayerNorm

As shown in Algorithm 1, we choose the pre-LayerNorm operation which places the layer normalization between the residual blocks. As suggested by recent studies, pre-LayerNorm yields more stable gradients especially at initialization (*27–30*). Current prevalent implementations of normalization layers in different deep learning packages usually contain element-wise affine transformations with learnable parameters immediately followed by the linear operations in many pre-layernorm Transformers including our designed architecture. This configuration is mathematically redundant except for minor differences caused by choice of optimizers during training. We therefore remove all element-wise affine transformations in the pre-LayerNorm.

**Figure S1:**
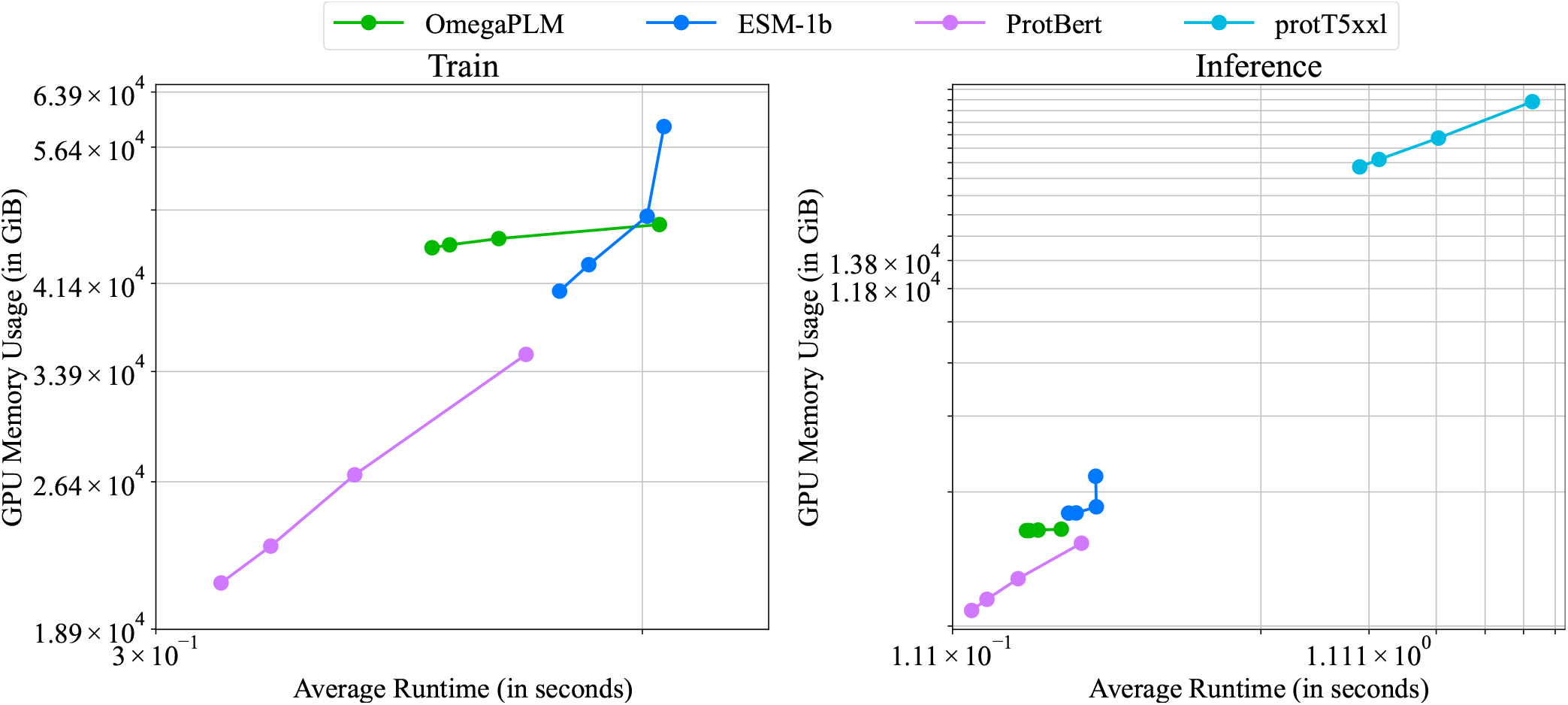
OmegaPLM strikes a great balance between efficiency and performance. This figure is in log scale both in time and in space. Each model in the plot has four points, where each one grows from 128 to 1024 exponentially in sequence length while decrease from 64 to 8 in batch size simultaneously. Points of all models in this plot go from bottom left to top right. ProtT5 cannot fit on our testing GPU (Nvidia A100 80GB) with the same data size during training. Value as mean from best 64 rounds of 128 rounds in total.

##### Gated Attention Unit

Instead of using multi-headed self-attention (MHSA), we adopt the Gated Attention Unit (GAU) (line 8 in Algorithm 1), which has shown great promise as an alternative to MHSAs with smaller memory consumption and faster convergence rate (*25*). We apply the gate operation after the attention aggregation and replace the conventional softmax(·) function with relu^2^(·) to aggregate the pairwise logits. In particular, we use an extra gating vector 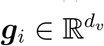 where d_*v*_ is the dimensionality of the value vector, which later multiply with the weighted summation of value ***v***_*j*_ in an element-wise fashion (line 8).

**Table S1:**
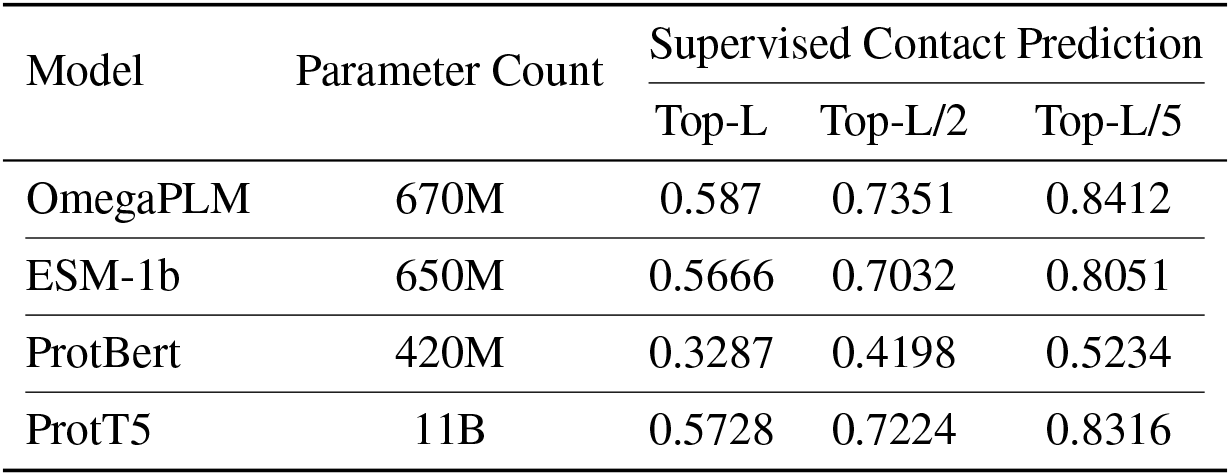
Supervised contact prediction performance shows OmegaPLM clears performs well. The contact prediction head is made of ResNets with identical configurations. The training (6,292 proteins) and test (699 proteins) datasets were selected as subsets of PDB (each protein having < 25% sequence similarity) using the following filtering conditions: (i) sequence length between 40 and 700, (ii) resolution better than 2.5 Å, (iii) no domains made up of multiple protein chains, (iv) PDB file with valid C_*β*_ coordinates (C_*α*_ in GLY) no less than 80% compare to the complete protein sequence. Our validation data keeps the same with (*26*) except 105 CASP11 test proteins. Proteins having sequence similarity > 25% or BLAST E-value < 0.1 with any test protein were excluded from training data.

| Model | Parameter Count | Supervised Contact Prediction |  |  |
| --- | --- | --- | --- | --- |
|  |  | Top-L | Top-L/2 | Top-L/5 |
| OmegaPLM | 670M | 0.587 | 0.7351 | 0.8412 |
| ESM-1b | 650M | 0.5666 | 0.7032 | 0.8051 |
| ProtBert | 420M | 0.3287 | 0.4198 | 0.5234 |
| ProtT5 | 11B | 0.5728 | 0.7224 | 0.8316 |

Similar to conventional transformer models, All the value vectors ***v***_*i*_, base vectors ***u***_*i*_ and gating vectors ***g***_*i*_ in GAU (*25*) are also generated by a sequence of Linear-SiLU operations, which later are used to produce queries and keys by using element-wise affine operations (line 4, line 5 & line 6 in Algorithm 1). The original GAU in (*25*) also suggests that relu^2^(·) performs better compared to softmax in terms of both computation speed and convergence rate. This performance and speed gain can only be achieved when the lengths of training and test proteins are relatively the same. However, it is common that the length of proteins to be predicted is out of the range of the lengths of training samples. To address this problem, we choose the softmax operation since it performs better than relu^2^(·) for sustaining the output distribution of the aggregated value vectors, as the summation of the attention weights is always 1. Empirically, we find that models trained with relu^2^(·) performs considerably worse than their counterparts with softmax when the input sequence length is out of the range of the trained samples, which is consistent with another study that models trained with long sequences perform subpar with shorter sequences when using relu^2^(·) (*31*).

There are a few other works targeting the extrapolation performance for the attention mechanism. (*32*) introduce a hand-craft relative positional bias term added to the pre-softmax logits. (*33*) argues for scaling the qk-product not only with inverse square root of d_*k*_, but also with log n, where n is the number of tokens in one sequence. When viewed as a distribution and under certain assumptions, attention scores’ entropy from the softmax operation oscillates less with varying sequence lengths with the log n scaling (*33*) compared to normal 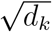 which improves generalization in terms of sequence length.

##### Relative Positional Encoding (RoPE)

Attention mechanism is inherently permutation-invariant so it requires position information when applied to sequence data. Here we apply rotary positional embedding (RoPE) (*34*) (line 5 & line 6 in Algorithm 1)) to encode the position information for a pair of amino acids, which is defined in Algorithm 2. (*34*) solved this question using the properties of complex numbers, and we apply such mechanisms into our queries and keys. To further emphasize the impact of relative position information, we include a bias term b_*i*−*j*_ which is specific to positions i and j. Note that the values of b_*i*−*j*_ and b_*j*−*i*_ are different. Instead of decreasing the embedding values as the absolute relative position increases (*32*), we clip the relative positions to allow extrapolation.

###### Algorithm 2

Rotary Relative Positional Embedding

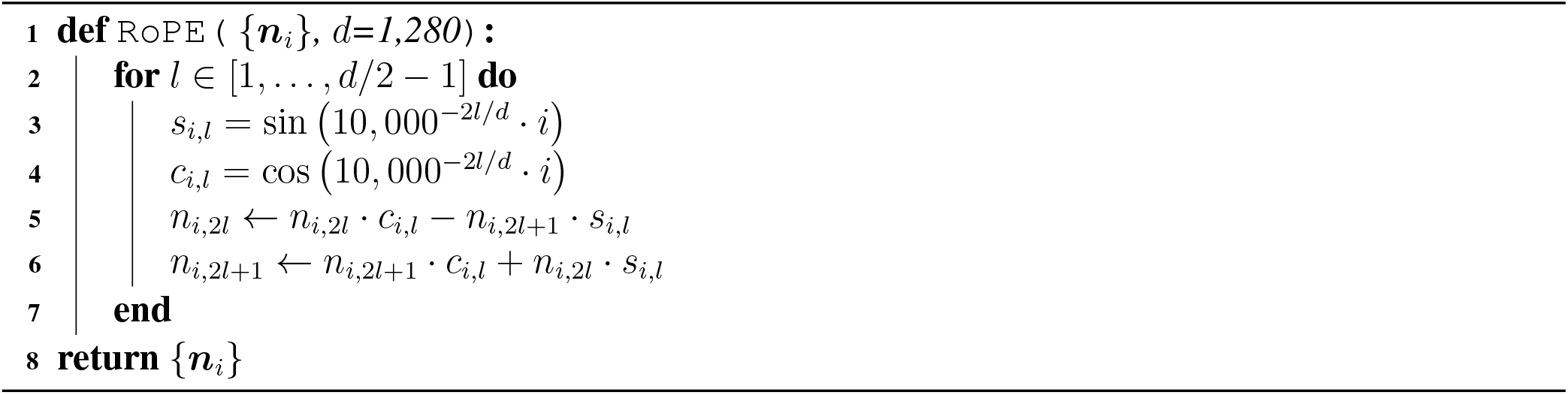

**Table S2:** OmegaPLM Configurations.

|  |  |
| --- | --- |
| No. Layers | 66 |
| $d$ | 1280 |
| $d_k$ | 256 |
| $d_v$ | 2560 |
| No. Attention Head | 1 |
| Tying Embeddings for input & prediction head | True |
| Cosine normalization with learned scale (28, 36) at output | True |
| Clipping thresholds of relative positional bias | [-64, 64] |
| Normalization type | LayerNorm (37) |
| Pre- or Post-Norm | Pre-Norm |
| Learnable parameters in normalization | False |

##### Hyper-parameters of OmegaPLM

The model configurations and the hyper-parameters are presented in Table S2. We generate the output tokens (*35*) using tied embeddings and cosine normalization (*28, 36*). The clipping thresholds in Table S2 indicate that the relative distance outside the range will be rounded to the thresholds.

#### 1.2 Training

##### Training Objective

Unlike ESM-1b (*12*) or any other protein language models (*38*), to train OmegaPLM, we setup a number of different objectives, we adopt the objective functions used in BERT (*1*) and spanBERT (*39*) as our framework upon which we modify different parts to acquire different variants of the objective function as follows,

1. BERT loss. For each sequence, 15% tokens are selected as targets to be predicted, 80% of which are replaced with a [mask] token, 10% of which are replaced with a random amino acid, and the final 10% stay what they are. This procedure is the same as the loss function used in the ESM-1b work (*12*);
2. SpanBERT-like loss. We sample the span length from Poisson distribution with λ = 7 and clip the sampled value at 5 and 8 and then mask the tokens consecutively according to the span length. Unlike SpanBERT, we still use the output embeddings from the corresponding tokens to perform prediction rather than the boundary tokens of the spans.
3. Sequential masking, where we mask either the first half or the second half of the sequence, akin to Prefix Language Modeling (*4, 40*)

Moreover, we assign different weights for these loss terms. The weights for the first two loss functions are 0.45 and the last one is 0.1.

##### Focal Loss

We observe that many of the amino acids can be accurately predicted with its short-range sequence context. This creates an easy prediction task for the model to learn and causes the model to overly focus on short-range relations. To address this problem, we adopt the focal loss (*41*) to down-weight the easy targets and make the model focus more on capturing the long range relationships among different amino acids.

##### Data

We train OmegaPLM using the Uniref50 (*42*) dataset with version 2021 04. During the training process, we randomly sample center protein sequences from Uiref50 (*42*) as the input data, where each batch contains 4,096 sequences and each sequence is padded or cropped to 512 residues. To fit one batch into the GPU memory, we adopt the gradient checkpoint technique on 23 of the 66 layers in the training process. Since OmegaPLM is trained for 3D structure prediction, we utilize 99.99% of the data for training and the rest of sequences for validation.

##### Parameter Initialization

All the weights in the output projection layers and biases are initialized as zeros following (*20, 43*). All the other weights are initialized using the normal distribution with zero mean and standard deviation of 0.02 following (*25*).

##### Recipe

OmegaPLM is implemented in PyTorch (*44*) and trained for 2,560 GPU Nvidia A100 80G days. We use the token-dropout scheme as in the ESM-1b model (*12*) during pre-training and AdamW (*45*) algorithm as our optimizer for OmegaPLM. The peak learning rate is set to 5e-4 and all the other parameters of AdamW stay as default in PyTorch. The warm-up of learning rate starts from 2.5e-6 and follows a cosine scheme for 12.5k steps, the linear weight decay starts at step 100k and lasts 500k steps. The learning rate stays constant at 2e-5 for another 100k steps. In addition, gradients of all parameters are clipped based on the norm at 0.3 (*46*) to address the spikes of the loss and gradient norms during training (*47–50*). In the hardware setup, we find the training process is accelerated by incorporating the PowerSGD (*51*) gradient compression with rank 32 to reduce the communication loads across different GPUs. Empirically we find PowerSGD improve 30% of the training speed. Though this compression introduces noise into the gradients, we find such noise inconsequential compared to the speed gain in convergence. Default precision format in Nvidia A100 GPUs is set to TensorFloat-32 for matrix operations. We do not change this setting as IEEE float16 causes overflow issues during training and IEEE float32 slows computation considerably. In validation, we clip the long proteins to the first 1,024 residues.

### 2 Geoformer: Geometric Transformer for Structure Prediction

#### 2.1 Geoformer Architecture

The 3D structure prediction of AlphaFold2 is driven by two important modules: triangle attention on edge representations and the Invariant Point Attention (IPA) in Structure Module. The triangle attention relies on the “triangle multiplicative update” on the edge representations to enforce the edge representation to satisfy the triangle inequality of amino-acid distances, which significantly improves the feasibility of predicted structures based on the MSA data. The geometric consistency of AlphaFold2 is maintained on the high-dimensional edge representations which later are integrated into node representations to generate the coordinate of each atom. However, geometric consistency in the high-dimensional space does not guarantee consistency in the Euclidean space because of the large amount of non-linear operations applied to the high-dimensional representation in the model. This posts a new computational challenge that on one hand the edge representations are to be optimized in the high-dimensional space to sufficiently capture the rich geometric information while on the other hand the real geometric consistency can only be checked after the representations are projected into the Euclidean space. To address this problem, we design a new structural module, named Geoformer, which could optimize high-dimensional representations to capture complex interaction patterns among amino acids while still maintaining geometric consistency in the Euclidean space.

##### Algorithm 3

Geometric Transformer

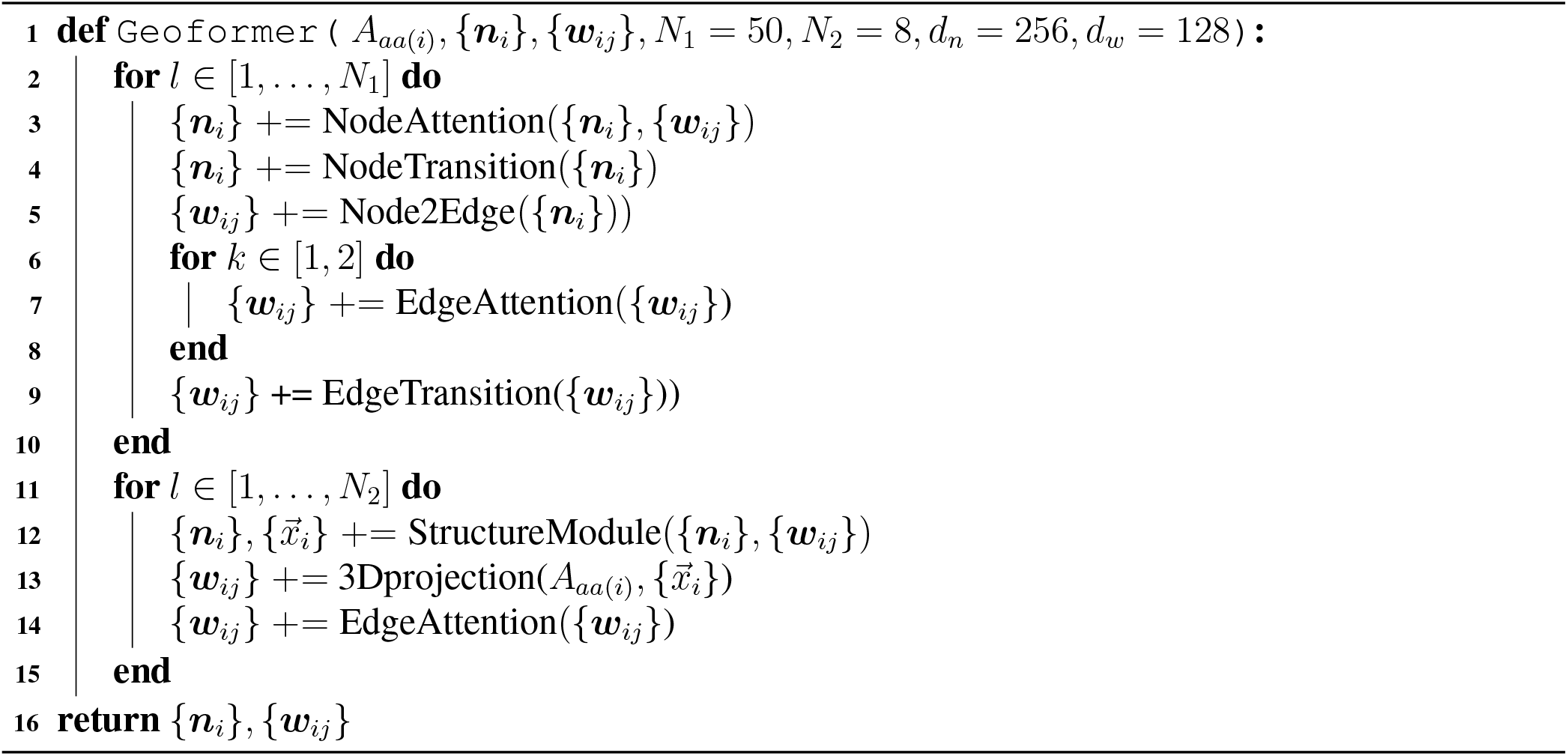

As shown in Algorithm 3, we update the node embedding {***n***_*i*_} based on two factors: 1) the attention between node *i* and any other node *j* and 2) the edge embedding {***w***_*ij*_} capturing the interactions between *i* and *j* in a more direct way. For each node, Geoformer first aggregates the information from all the other nodes to generate a basic node representation (*NodeAttention* and *NodeTransition* functions in line 3-4 in Algorithm 3). Next, we produce another temporal edge representation solely based on the node representation inferred from the previous step using the function *Node2Edge*. Similar to *NodeAttention*, we also update edge representations {***w***_*ij*_} based on all the other edges using a transformer-based model *EdgeAttention*, which is also the function we rely on to achieve geometric consistency in the high dimensional space. We repeat this process 50 times to generate both node and edge representations for each residue and residue pair. In the last 8 layers of Geoformer (lines 11-15), we aim to maintain the geometric consistency in the Euclidean space. In particular, we first translate the inferred node and edge representation of a protein to a temporal 3D structure by using the *StructureModule* function implemented by AlphaFold2 (line 24-31 in Algorithm 20 in AlphaFold2). x_*i*_ is the coordinates for the atoms in amino acid *i*. We then translate the temporal 3D structure back to the high dimensional space by using the 3*Dprojection* function whose outputs have the same dimensionality as the ***w***_*ij*_. In this way, the updated edge representation contains the information indirectly encoded from the 3D space. Another *EdgeAttention* function is applied to achieve geometric consistency for the newly updated ***w***_*ij*_. Eventually, both node representation ***n***_*i*_ and edge representation ***w***_*ij*_ from the last layer are used to predict the 3D coordinates and connected to the loss functions. Note that we omit the LayerNorm operation on the set variables such as {***w***_*ij*_} to simplify the notation.

#### 2.2 Node Update

The NodeAttention function provides expressive node representations by integrating all the node and edge representations using the self-attention layers. The computational graph is defined as the follows,

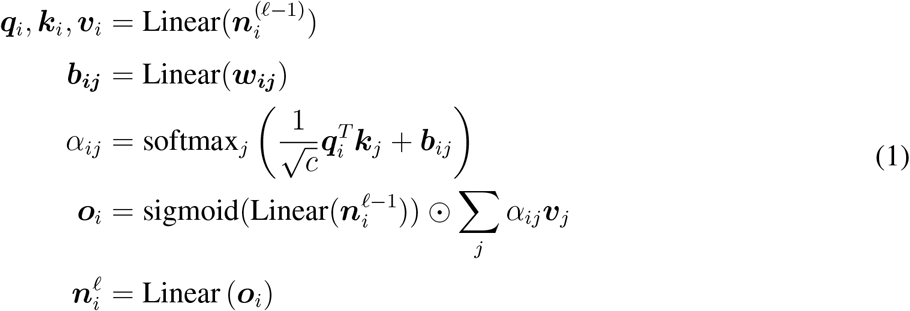

Here ***n***_*i*_ and ***w***_*ij*_ in the first layer are the node and edge representations extracted from the OmegaPLM module, respectively. ***b***_***ij***_ is a linear projection of ***w***_*ij*_, which is used as the bias term inside the attention function capturing the interactions between node embedding ***n***_***i***_ and ***n***_***j***_. Similar to the Algorithm 1, we also find the presence of the gate operation allows a much simpler attention mechanism than conventional MHSA without too much quality loss. *NodeTransition* in Algorithm 3 is a standard two feed-forward layer with *relu* activation function.

#### 2.3 Edge Update

*EdgeAttention* achieves geometric consistency by simultaneously considering all the other edge embeddings ***w***_*ij*_ using the following equations,

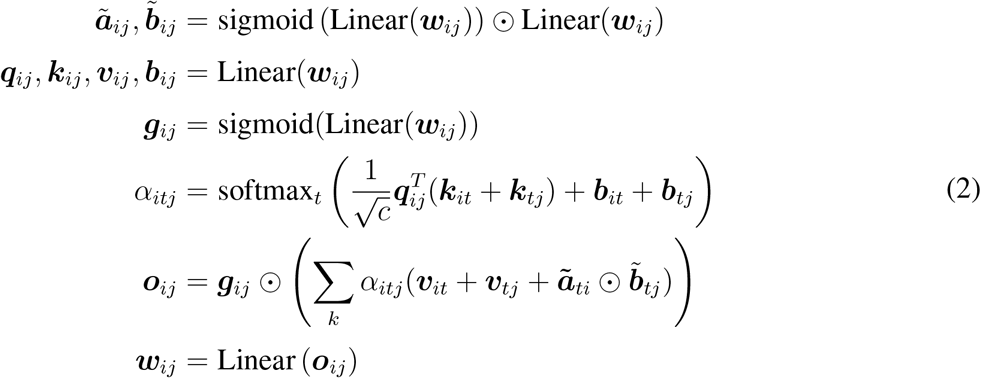

The edge embedding ***w***_*ij*_ is updated based on various types of interactions involving a third node *t*. Note that AlphaFold2 uses four triangular multiplicative operations to maintain this consistency including “outgoing edge attention”, “incoming edge attention”, “starting node attention”, and “ending node attention”. As shown in Equation 2, *EdgeAttention* defines two new interaction terms which integrate both the self-attention and bi-linear models. First, instead of using different self-attention mechanisms as AlphaFold2, we add the key value ***k***_***it***_ and ***k***_***tj***_ together as the new key value and multiply with query ***q***_***ij***_. So far as we know, this is the first model which leverages the summation of the key values to capture the interactions among three entities. Second, we integrate the bi-linear interaction between ***a***_***ti***_ and ***b***_***tj***_ is inside the attention function for calculating ***o***_***ij***_. In this way, we design one universal function to implement the four functions of AlphaFold2, which significantly reduces the computational burden. Similar to *NodeTransition, EdgeTransition* is a standard two feed-forward layer with the relu activation function. In the *Node2Edge* function, edge embedding ***w***_*ij*_ is updated by the outer product of linear projection of ***n***_***i***_ and ***n***_***j***_.

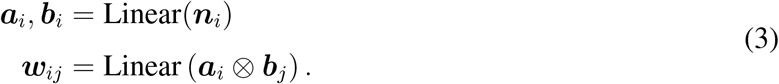

#### 2.4 Geometric Atom Embedder

The last eight layer of Geoformer includes the Geometric Atom Embedder, where we incorporate the Invariant Point Attention (IPA) module and the rotation/transition matrix representations in AlphaFold2 for the coordinate generation (line 12-14 in Algorithm 3). The *StructureModule* function translates the node embedding ***n***_*i*_ to a protein structure, which is represented by the 3D coordinates 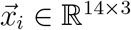 for amino acid i. An amino acid has a maximum of 14 atoms and each atom is associated with a 3D coordinate. Function *StructureModule* (line 24-31 in Algorithm 20 in AlphaFold2) predicts 3D coordinates by optimizing the Transition and Rotation matrices. Based on this temporal 3D structure, function 3*Dprojection* produces a new type of interaction based on three types of information extracted from the predicted temporal protein structures {***x***_*i*_}: 1) the amino acid types of *i* and *j*; 2) the relative positions of each atom in amino acid i and j in different frames; 3) the relative angles between the vectors of each atom in amino acid *i* and *j* in different frames. The 3D-based interaction vector is projected by a Multilayer Perceptron (MLP) to match the dimension of edge embedding. The function 3*Dprojection* is defined in Algorithm 4.

##### Algorithm 4

3Dprojection

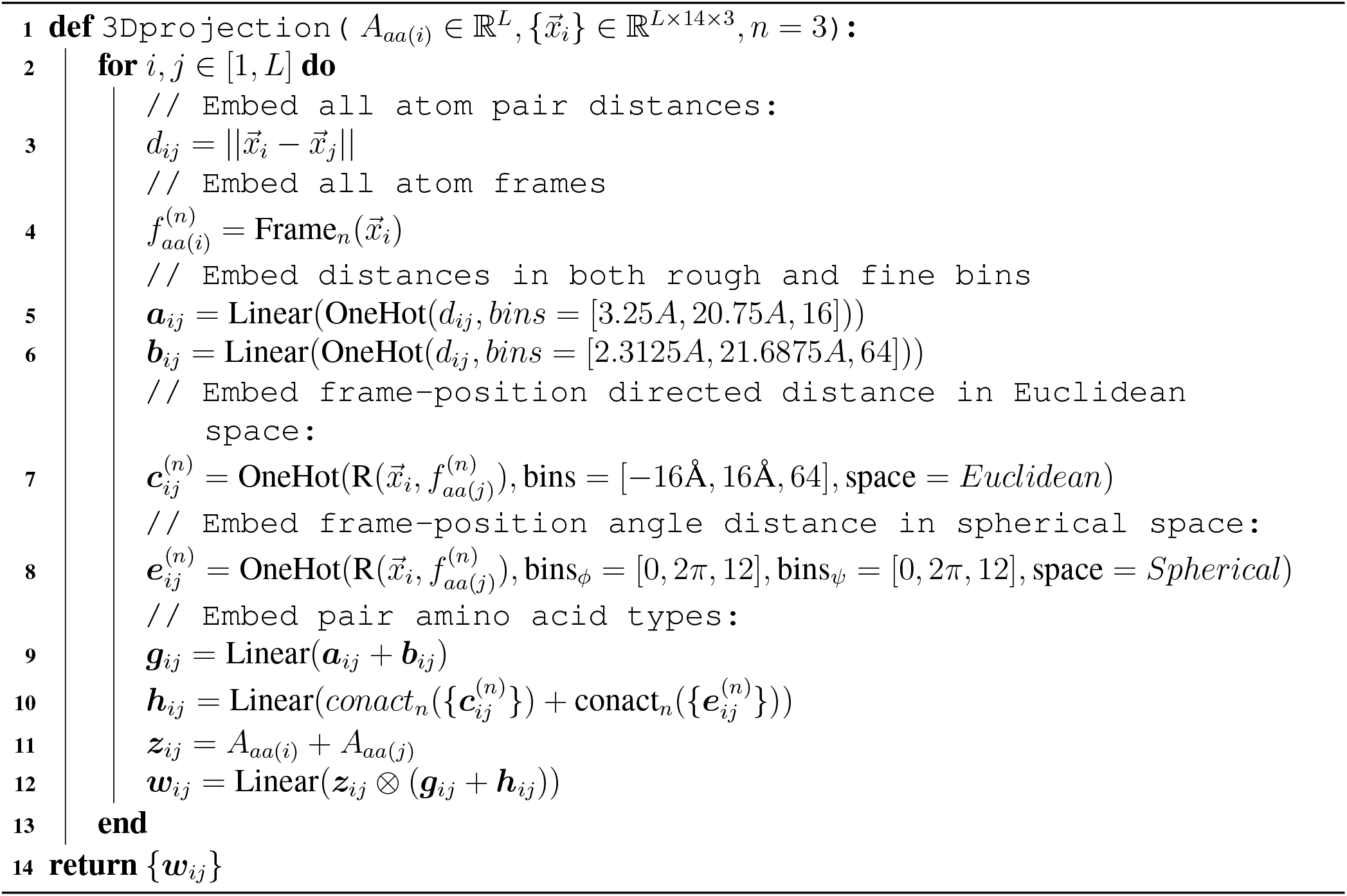

Here *d*_*ij*_ is the distance between atoms *i* and *j* from two amino acids. ***a***_*ij*_ and ***b***_*ij*_ are embeddings based on one hot vectors of Euclidean distance between atoms *i* and *j*. The difference between ***a***_*ij*_ and ***b***_*ij*_ is the ways of their bin constructions. Function Frame_*n*_ returns the frame 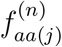 that represents the local coordinate system with *C*_*α*_, *C*_*β*_ and N atoms of amino acid *aa*(*j*), and function **R** is a projection function which returns the 3D coordinates *x*_*i*_ in the *n*-th coordinate systems 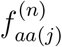 defined based on the amino acid associated with atom *j*. For each amino acid, we use eight-frame coordinate systems defined by AlphaFold2 (Section 1.8 in the supplementary of AlphaFold2). *aa*(*i*) is the amino acid type that atom i belongs to. ***z***_***ij***_ represents the one-hot embedding related to the types of amino acids that atoms *i* and *j* are associated with. We use an embedding matrix *A* ∈ ℝ^20×*d*^ to represent each of the 20 types of amino acids, where *d* denotes the dimension of the continuous embedding. Note that this embedding is not position-specific and we refer to this embedding as *type embedding* because it corresponds to the type of amino acids. ***c***_*ij*_ and ***e***_*ij*_ are designed to model the relative positions of two atoms in different coordinate systems in both the Euclidean space and the spherical space. In particular, as it requires two angles *ϕ* and *ψ* to describe a location in the spherical space, we discretize these two angles into two sets of bins bin_*ϕ*_ and bin_*ψ*_. The amino-acid embedding ***z***_*ij*_ and physical location embeddings ***g***_*ij*_ and ***h***_*ij*_ are later integrated together by using the outer product operation.

**Figure S2:**
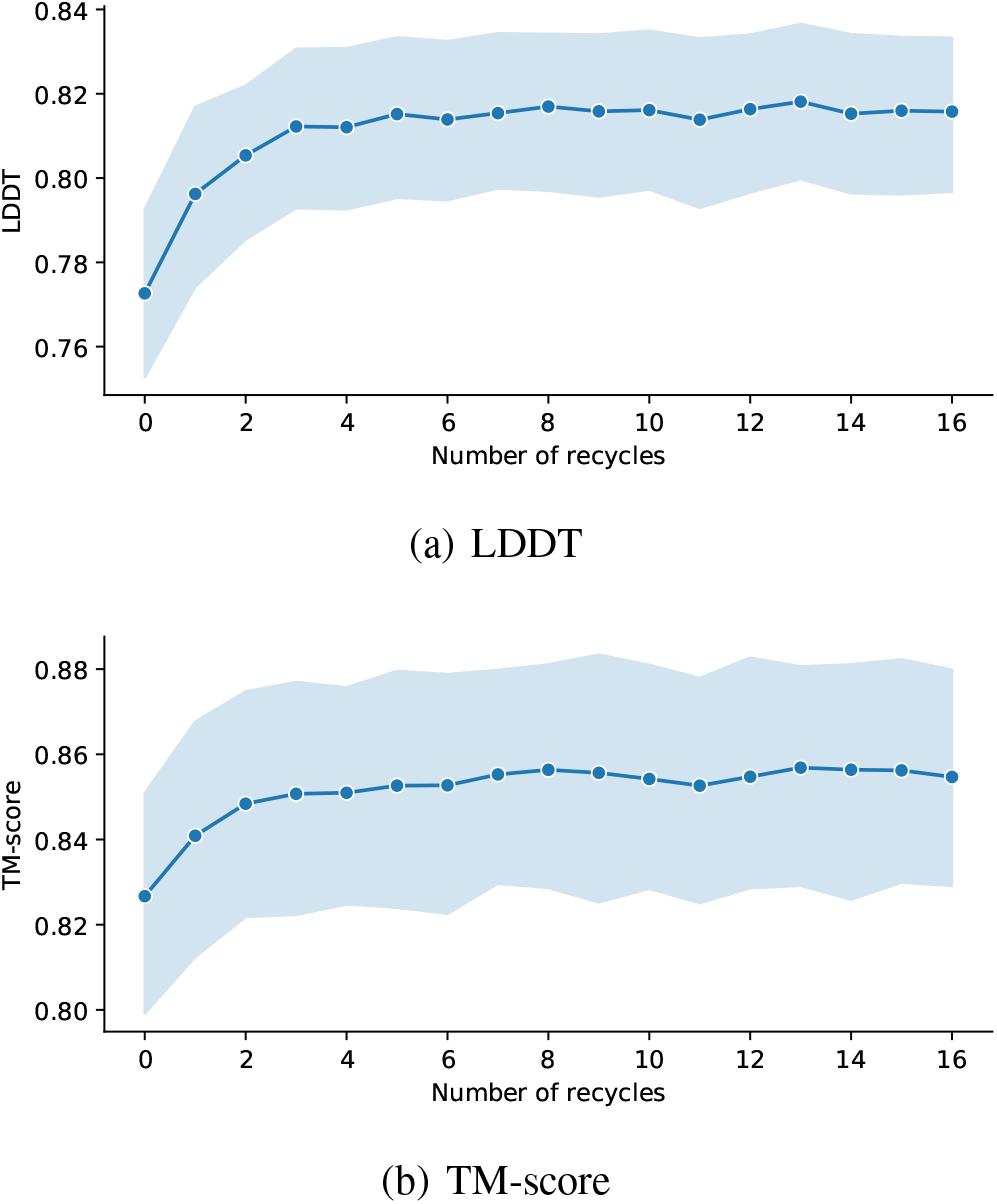
Performance of structure prediction with recycling. OmegaFold reaches reasonable performance without recycling, and achieves highest performance after around 10 recycles.

#### 2.5 Recycling

Similar to AlphaFold2 and RoseTTAFold, we also employ a recycling procedure to iteratively refine the quality of structure prediction. We find that with the prediction accuracy improves with the number of recycles and saturates after about 10 recycles (**Fig. S2**).

#### 2.6 Datasets

We download the protein structure data from the PDB website (https://www.rcsb.org/) and split each PDB file into multiple single chains. A protein is removed if 1) it contains RNA fragments or 2) more than 90% of its residues are unknown amino acids which are marked as ‘X’ or 3) one of its chains can not be parsed by PDBparser of biopython (*52*). Next, we cluster all the sequences using MMseqs2 (*53*) with sequence identity threshold 100% and only keep the cluster centers. For the selected proteins, we perform 40% clustering using MMseqs2 and calculate a static sampling rate for each protein chain. The rate is inverse to the size of the cluster the sample belongs to.

##### 2.6.1 Validation Data

We use two validation datasets. The first CASP dataset includes challenging free-modeling (FM) targets from recent CASP13 and CASP14: T0950, T0953s1, T0953s2, T0955, T0957s1, T0957s2, T0958, T0960, T0963, T0968s1, T0968s2, T1005, T1008, T1026, T1027, T1029, T1030, T1031, T1032, T1033, T1038, T1043, T1046s1, T1046s2, T1049, T1056, T1064, T1082, T1099.

A second dataset includes recently targets with a wide range of prediction difficulty from CAMEO from date1 to date2: 7RPY:A, 7PB4:I, 7FJS:L, 7U2R:A, 7AAL:A, 7KO9:A, 7RKC:A, 7EQH:A, 7U4H:A, 7ETS:B, 7E4S:A, 7F3A:A, 7LXK:A, 7EAD:A, 7DKI:A, 7ESH:A, 7VNO:A, 7Z79:B, 7RW4:A, 7W5M:A, 7WNW:B, 7OPB:D, 7EQE:A, 7N0E:A, 7T4Z:A, 7ED6:A, 7NUV:A, 7TV9:C, 7ZCL:B, 7VWT:A, 7PC6:A, 7NQD:B, 7TXP:A, 7PC4:A, 7QRY:B, 7FEV:A, 7FIW:B, 7RPR:A, 7OA7:A, 7EBQ:A, 7YWG:B, 7UGH:A, 7F0A:A, 7U5Y:A, 7NDE:A, 7QIL:A, 7X4E:A, 7OD9:C, 7TBU:A, 7W26:A, 7X4O:B, 7NMI:B, 7WRK:A, 7QSS:A, 7LI0:A, 7RGV:A, 7VSP:C, 7X8J:A, 7QDV:A, 7E52:A, 7RCW:A, 7TNI:C, 7PC3:A, 7N29:B, 7F2Y:A, 7ZGF:A, 7T03:A, 7MYV:B, 7BLL:A, 7MQ4:A, 7X9E:A, 7F6J:C, 7EJG:C, 7V4S:A, 7QAP:A, 7ACY:B, 7MLA:B, 7QAO:A, 7WWR:A, 7QSU:A, 7PZJ:A, 7V1K:A, 7SGN:C, 7Z5P:A, 7N3T:C, 7EGT:B, 7O4O:A, 7CTX:B, 7VNX:A, 7YXG:A, 7QS5:A, 7X8V:A, 7MKU:A, 7RPS:A, 7MS2:A, 7QBP:A, 7QS2:A, 7EHG:E, 7PRQ:B, 7S2R:B, 7R74:B, 7W5U:A, 7O0B:A, 7TA5:A, 7WWX:A, 7Q51:A, 7SXB:A, 7WCJ:A, 7LXS:A, 7OSW:A, 7WRP:A, 7PNO:D, 7WJ9:A, 7RCZ:A, 7U5F:D, 7WME:A, 7RXE:A, 7B0K:A, 7ERN:C, 7R63:C, 7SJL:A, 7BI4:A, 7W1F:B, 7ED1:A, 7RAW:A, 7SO5:H, 7VOH:A, 7Q05:E, 7QBZ:A, 7PSG:C, 7P0H:A, 7MHW:A, 7P3I:B, 7ULH:A, 7R09:A, 7F0O:B, 7EQB:A, 7EFS:D, 7TZE:C, 7W74:A, 7PXY:A, 7PW1:A, 7E5J:A, 7V8E:B, 7ERP:B, 7R5Y:E.

#### 2.7 Structure-based Sampling

Given limited computation resources, we need to crop long proteins to fit the GPU memory for each learning epoch. Previous deep learning models usually sample a sub-sequence with fixed length (*20*). Adjacent residues in a protein sequence are usually closed in the 3D structure. However, sampling and cropping proteins solely based on protein sequences will decrease the number of residue pairs in which two amino acids are far apart in the sequence but close in the 3D space. To address this problem, we introduce the consecutive structure sampling approach to allow the model to extract long-range contact information from the data pool more efficiently. We first cluster all the sequences at the chain level. Each cluster and each protein in the cluster have the same probability to be sampled. Within each sampled protein, we uniformly sample a residue *r*_*c*_ and then sample a continuous structural fragment centered around this by maximizing the following function,

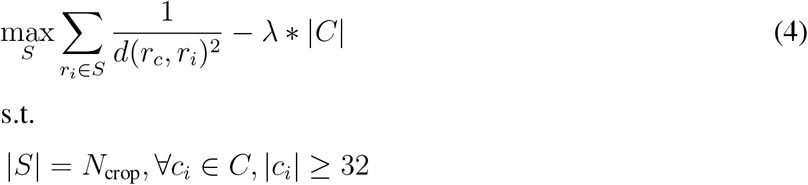

where function *d*(·, ·) measures the Euclidean distance between the *C*_*β*_ atoms of a pair of amino acids (for Alanine we use its *C*_*α*_ atom). *S* is the residue set we already select and N_*crop*_ is total number of the amino acids we want to select based on the size of GPU memory. *C* is the set of continuous sub-sequences in this select 3D structure fragment, |*C*| is the number of sub-sequences and λ is a pre-defined hyper-parameter which is set as 0.004 in experiments. For each sub-sequence *c*_*i*_ ∈ *C*, the residues are continuous on the sequence and located on the same chain. The central idea of this optimization problem is that we want to select *N*_*crop*_ amino acids around *r*_*c*_ in the 3D space without too many sub-sequences, which is implemented by adding the second term to penalize the number of sub-sequences. At the same time, we also require the length of each sub-sequence *c*_*i*_ to be at least 32 to prohibit too short sub-sequences.

Equation 4 could be solved by dynamic programming in a very efficient way. For a protein to be cropped, we first concatenate all the chains of that protein into one sequence with n chains and a total length of L. Then we design a temporal variable *F*_*i,j*_(0 ≤ *i* ≤ *L*, 0 ≤ *j* ≤ *N*_*crop*_), which indicates that the maximum score we could achieve among the first *i* residues. At this step, we select *j* residues and residue *i* has to be one of them. All the sub-sequences have at least length 32. We use another temporal variable *G*_*i,j*_(0 ≤ *i* ≤ *L*, 0 ≤ *j* ≤ *N*_*crop*_) to denote the maximum value among the first i residues. However, residue i is not necessary in the j residues we select. We need to require all the fragments to have at least length 32. The relationship between *F*_*i*_ and *G*_*i*_ is *F*_*i*_ = max_*j*∈[0,*i*]_ *G*_*j*_. The only difference between *F* and *G* is that F requires residue *i* in the selected residue set but *G* does not have this requirement. *F* and *G* can be calculated by using the following recursion formula,

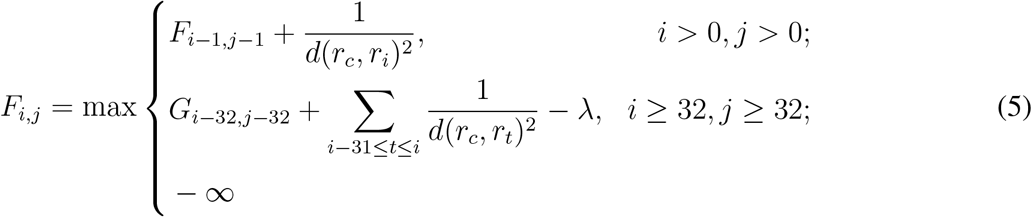

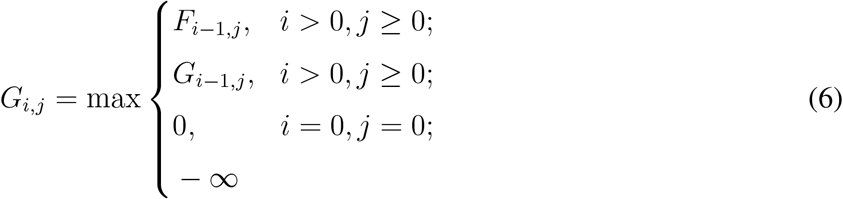

**Table S3:** OmegaFold Training Protocol

| Model | First Stage | Second Stage | Last Stage |
| --- | --- | --- | --- |
| Sequence crop size | 256 | 320 | 384 |
| Batch size | 128 | 128 | 128 |
| Learning rate | 5e-4 | 1e-4 | 5e-5 |
| Structural violation loss weight | 0.0 | 1.0 | 1.0 |
| Dynamic sampling strategy | Average Sampling | Hard Sample Mining | Gradient Harmonization |
| Training time | $\approx 7$d | $\approx 10$d | $\approx 10$d |
| Number of epoches | 20k | 20k | 15k |

The maximum value of Equation 4 is obtained at 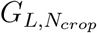. By recording the generation process of this optimal solution, we can collect the amino acids on the path to get sets S and C. For a selected set S, we first generate the sequence embedding for each sub-sequence using OmegaPLM separately and then concatenate them together as the embedding of the 3D structure fragment. In order to use OmegaPLM, we need to specify a new residue index to calculate the positional embedding as the new data is not continuous. We set the gap as 64 for the residue index between different sub-sequences, making the pair embedding of different sub-sequences into two separate bins.

##### Curriculum Learning with Dynamic Sampling

In the training process of OmegaFold, we dynamically change the sampling rate of each training data because we find that there exists a significant amount of protein chains whose structures are hard to predict. One of the reasons is that a protein chain might need to interact with other biological entities such as RNAs and proteins to form a stable 3D structure. There is another set of proteins for which 3D structures are inherently hard to predict as there are few homologous templates existing in the training data or the co-evolution information extracted from data is sparse.

To alleviate the influence of proteins without stable structures and encourage the learning of hard proteins, we assign each protein 3D fragment a sampling rate and adjust the sampling rate according to the LDDT score for the Cα atom calculated based on the temporal predicted structures during training. If the fragment has never been calculated by the model, the sampling rate is set to 1.0. If only a part of the fragment has been trained, the sampling rate is adjusted according to the partial prediction. In particular, we adopt three different strategies for three different training stages. At the first training stage (20,000 epochs), we do not add extra sampling rate adjustment to stabilize the training process. At the second training stage (20,000 epochs), we lower the sampling rate of fragments with an average per-residue LDDT-Cα greater than 0.6. The sampling rate factor linearly decreases from 1.0 to 0.0 when the LDDT-Cα ranges from 0.6 to 1.0, which encourages the model to focus on hard samples. At the last training stage (15,000 epochs), inspired by (*54*), we lower the sampling rate of fragments with the average LDDT-Cα either greater than 0.9 or less than 0.5, which indicates they are either easy or unstable proteins that are impossible to fit. Specifically, when the LDDT-Cα ranges from 0.0 to 0.5, the sampling rate linearly increases from 0.5 to 1.0. When the LDDT-C*α* is between 0.5 and 0.9, the sampling rates are all set to 1.0. When the LDDT-Cα ranges from 0.9 to 1.0, the sampling rate decreases from 1.0 to 0.0.

**Table S4:** Ablation Study

| Model | validation LDDT |
| --- | --- |
| OmegaFold (ensemble) | 0.816 |
| OmegaFold (single) | 0.801 |
| AlphaFold2 with single sequence input | 0.344 |
| Retrained AlphaFold2 with single sequence input | 0.476 |
| OmegaPLM with structure module (no Geoformer) | 0.198 |

#### 2.8 Training

##### Loss function

We train the OmegaFold system using the contact loss, FAPE loss, torsion angle loss, and structure violation loss as in AlphaFold2. We discard experimentally resolved loss which does not influence the folding performance. We also discard masked MSA prediction loss since we no longer use MSAs as input.

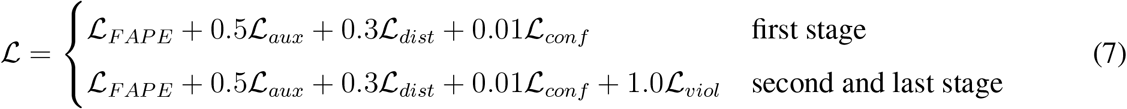

##### Parameter Initialization

The weights of all the output linear layers are initialized as zeroes and the biases in linear layers before the sigmoid function are initialized as ones while the weights are zeros. All the other linear layers are initialized with Glorot Uniform initialization.

##### Recipe

The protein language model and 3D structure prediction model are trained separately as the protein language model includes more unlabeled training data. In this stage, we freeze the protein language model and only optimize the 3D structure prediction model by using the AdamW (*55*) algorithm with default parameters. The batch size stays 128 through the whole training process. The learning rate linearly increases from 0 to 5e-4 during the first 1,000 steps, stays 5e-4 for next 19,000 steps, then changes to 1e-4 for another 20,000 steps and is fixed as 5e-5 for the last 15,000 steps (Table S3). We also clip the gradient norm using the threshold of 0.1. We use the function torch.nn.utils.clip grad norm in PyTorch (*56*), which clips the gradient after collecting the gradients of all the samples using the all-reduce operation.

#### 2.9 Ablation Study

We conduct a series of ablation studies on the OmegaFold system. To evaluate the contribution of OmegaPLM and Geoformer, we test models with the following configurations on the CAMEO validation dataset: the original ensemble version of OmegaFold, a single OmegaFold model without ensembling, DeepMind’s AlphaFold2 with a single sequence as input, re-trained AlphaFold2 model with single-sequence input, and an end-to-end model with only OmegaPLM and structure module. **Table S4** indicates that both OmegaPLM and Geoformer have a critical contribution to OmegaFold’s superior performance. Notably, after retraining with single sequences as input, without the language model, AlphaFold2 is not able to predict structure with comparable performance.

## Notes

### Competing Interest Statement

The authors have declared no competing interest.

